# Association between SARS-CoV-2 and metagenomic content of samples from the Huanan Seafood Market

**DOI:** 10.1101/2023.04.25.538336

**Authors:** Jesse D. Bloom

## Abstract

The role of the Huanan Seafood Market in the early SARS-CoV-2 outbreak remains unclear. Recently the Chinese CDC released data from deep sequencing of environmental samples collected from the market after it was closed on January-1-2020 (Liu et al. 2023a). Prior to this release, Crits-Christoph *et al*. (2023) analyzed data from a subset of the samples. Both studies concurred that the samples contained genetic material from a variety of species, including some like raccoon dogs that are susceptible to SARS-CoV-2. However, neither study systematically analyzed the relationship between the amount of genetic material from SARS-CoV-2 and different animal species. Here I implement a fully reproducible computational pipeline that jointly analyzes the number of reads mapping to SARS-CoV-2 and the mitochondrial genomes of chordate species across the full set of samples. I validate the presence of genetic material from numerous species, and calculate mammalian mitochondrial compositions similar to those reported by Crits-Christoph *et al*. (2023). However, the number of SARS-CoV-2 reads is not consistently correlated with reads mapping to non-human susceptible species. For instance, 14 samples have >20% of their chordate mitochondrial material from raccoon dogs, but only one of these samples contains any SARS-CoV-2 reads, and that sample only has 1 of *∼*200,000,000 reads mapping to SARS-CoV-2. Instead, SARS-CoV-2 reads are most correlated with reads mapping to various fish, such as catfish and largemouth bass. These results suggest that while metagenomic analysis of the environmental samples is useful for identifying animals or animal products sold at the market, co-mingling of animal and viral genetic material is unlikely to reliably indicate whether any animals were infected by SARS-CoV-2.

---

Initial reports from Chinese officials about the outbreak that eventually became the SARS-CoV-2 pandemic described patients associated with the Huanan Seafood Market, and said there was no evidence of significant human-to-human transmission (ProMED 2019; Wuhan Municipal Health Commission 2019, 2020; WHO 2020). These two claims implied that the infections likely originated from a non-human source within the market.

But by mid to late January of 2020, it was clear SARS-CoV-2 was spreading from human to human, and had been for some time (Chan et al. 2020; Phan et al. 2020; Li et al. 2020; Nishiura *et al*. 2020). In addition, Chinese scientists published papers reporting that some of the earliest identified human cases in December 2019 had no link to the Huanan Seafood Market (Huang *et al*. 2020; Chen et al. 2020). Thus began a debate that continues to this day: was the market an initial source of zoonotic infections from animals, or was it simply a superspreading site that amplified earlier human infections from another source (Cohen 2020)?

It is universally agreed that there were human cases of SARSCoV-2 at the Huanan Seafood Market by mid December of 2019 (Li et al. 2020). It is also known that some animals susceptible to SARS-CoV-2 were sold at the market (Xiao et al. 2021). The unanswered question is if any animals were infected, and if so whether they infected humans or were infected by them.

In early 2022, the Chinese CDC posted a preprint on *Research Square* describing their sampling of the market beginning immediately after its closure on January-1-2020 (Liu et al. 2022). The Chinese CDC collected samples from both the environment and animals / animal products. They reported that none of the 457 animal samples tested positive for SARS-CoV-2, but that 73 of 923 environmental samples tested positive. A significant caveat of the Chinese CDC study is that all the samples were collected on January-1-2020 or later, which is *at least* a month after the first human infections in Wuhan (ODNI 2022; Zhang et al. 2020; van Dorp et al. 2020; Pipes et al. 2021). Despite this significant caveat, the data reported in the Chinese CDC study has been variously interpreted to support arguments that virus in the market was human derived (Liu et al. 2022), that the outbreak originated from live animals sold in the market (Worobey et al. 2022), or that the virus spread among humans in Mahjong rooms or toilets in the market (Courtier-Orgogozo and de Ribera 2022).

One aspect of the data gathered by the Chinese CDC that was *not* thoroughly analyzed in their 2022 preprint was the metagenomic content of the environmental samples. The 2022 Chinese CDC preprint simply reported that the number of SARSCoV-2 reads in deep sequencing of the samples was correlated with the number of human reads (fourth figure of Liu *et al*. (2022)), but did not specify correlations for other species and did not provide the raw sequencing data. Other scientists pointed out that if the raw data were shared, it would be possible to expand upon the analysis in the 2022 Chinese CDC preprint to determine if the abundance of SARS-CoV-2 genetic material correlated with material from other species (Cohen 2022a,b).

At some point after posting their 2022 preprint, the Chinese CDC uploaded raw sequencing data for some environmental samples to the GISAID database, where they were subsequently downloaded and analyzed by another group of scientists. News of this analysis leaked to the media, which published stories emphasizing the co-mingling of genetic material from raccoon dogs and SARS-CoV-2 in one of the samples (Wu 2023; Mueller 2023). The next week, the scientists published their report (Crits-Christoph et al. 2023), which described bioinformatic analyses showing that some environmental samples contained genetic material from animals susceptible to SARS-CoV-2, including raccoon dogs. However, although Crits-Christoph *et al*. (2023) analyzed the mammalian metagenomic content of the partial set of samples they obtained from GISAID, they did not report any analysis of the SARS-CoV-2 content of the samples.

A week later, the Chinese CDC posted an updated version of their preprint on the *ChinaXiv* server (Liu et al. 2023b) and released the full set of raw sequencing data in public databases. This updated preprint, and a version published by *Nature* the next week (Liu et al. 2023a) emphasized different aspects of the data than Crits-Christoph *et al*. (2023). Specifically, although the Chinese CDC concurred that some of the environmental samples contained material from raccoon dogs and other susceptible species, they stated that material from many species was found, and that raccoon dog material was more commonly identified in SARS-CoV-2 negative samples. However, like Crits-Christoph *et al*. (2023), the new version of the Chinese CDC paper (Liu et al. 2023a) did not analyze the association between the amount of SARS-CoV-2 and animal material in the samples, and even removed the figure from the original preprint (Liu et al. 2022) that showed that the number of SARS-CoV-2 reads was correlated with reads from both humans and several unidentified species.

Here I systematically analyze the relationship between the metagenomic and SARS-CoV-2 content of all environmental samples for which deep sequencing data are available. This analysis confirms that samples contain genetic material from a wide range of species, including some that are susceptible to SARS-CoV-2. However, the samples that contain abundant material from raccoon dogs and other non-human susceptible species usually contain little or no SARS-CoV-2 reads. Furthermore, the number of SARS-CoV-2 reads is not consistently correlated with reads mapping to non-human susceptible species, but is instead generally most correlated with reads from species that are implausible candidates for having been infected with SARSCoV-2. These results suggest that SARS-CoV-2 was widespread at the market by January 2020, and therefore that co-mingling of viral and animal genetic material in environmental samples collected at that time is unlikely to be informative about the original source of the outbreak.

## Results

### Deep sequencing data deposited in public databases on March-29-2023 by the Chinese CDC are a superset of the data analyzed by Crits-Christoph et al earlier that month

On March-29-2023, the Chinese CDC released raw deep sequencing data for samples taken from the Huanan Market in early 2020. These data were released on the NGDC database under accession CRA010170 (https://ngdc.cncb.ac.cn/gsa/browse/CRA010170).

I created a fully reproducible computational pipeline that downloaded and analyzed the full dataset from the NGDC (my pipeline is available at https://github.com/jbloom/Huanan_market_samples). The dataset consists of 696 FASTQ files from 395 deep sequencing runs of 176 samples, with the total size of the gzipped files exceeding three terabytes.

Prior to the full posting of the dataset on the NGDC by the Chinese CDC, Crits-Christoph *et al*. (2023) downloaded and analyzed 227 FASTQ files that had been uploaded to the GISAID database by the Chinese CDC. When news of the analysis by Crits-Christoph *et al*. (2023) became public, GISAID reportedly revoked access to the data and stated that the set of files that had been downloaded was incomplete (Cohen 2023a; GISAID 2023).

To determine the relationship between the 696 FASTQ files released by the Chinese CDC on the NGDC on March-29-2023 and the 227 files downloaded earlier that month by Crits-Christoph *et al*. (2023), I computed the SHA-512 hashes for all the NGDC FASTQ files and compared them to the hashes reported in Table S1 of Crits-Christoph *et al*. (2023). This analysis confirmed that the dataset released on the NGDC contained unmodified versions of all 227 FASTQ files downloaded by Crits-Christoph *et al*. (2023), plus an additional 469 FASTQ files (Tables S1 and S2).

### Analysis of mitochondrial sequences validates mammalian metagenomics by Crits-Christoph et al, and also finds abundant non-mammalian chordates

To determine the metagenomic content of the samples, I performed an analysis that conceptually parallels that described by Crits-Christoph *et al*. (2023). Throughout this paper, I report results only for sequencing described by the Chinese CDC as “RNA sequencing of total nucleic acids from environmental swabs for metagenomics,” and exclude the small number of samples that involve amplicon-based sequencing of SARS-CoV-2 or RNA sequencing for viral whole-genome assembly (Table S2).

Briefly, the deep sequencing data were pre-processed to retain only high-quality reads. These reads were aligned to the SARS-CoV-2 genome and a representative set of chordate mitochondrial genomes. This set included all the mammalian mitochondrial genomes from all species for which alignments were reported in Crits-Christoph *et al*. (2023). The accessions for the 4,170 chordate mitochondrial genomes that formed this set are listed in Table S3. The counts of reads mapping to each mitochondrial genome for each sequencing run and sample are provided in Tables S4 and S5, respectively.

I compared the mitochondrial compositions of the samples determined by my analysis to those reported by Crits-Christoph *et al*. (2023). Note that while Crits-Christoph *et al*. (2023) describe aligning the sequencing data to all metazoa mitochondria, they only report alignment counts for mammals. The aligned read counts for mammals from my analysis are highly correlated with those reported by Crits-Christoph *et al*. (2023) for all sequencing runs with a reasonable number of total counts (Figure 1). My analysis and that of Crits-Christoph *et al*. (2023) are independently implemented, and use different alignment programs, reference genome sets, and quality filtering criteria—so the fact that the two analyses give highly correlated results is reassuring.

**Figure 1.**
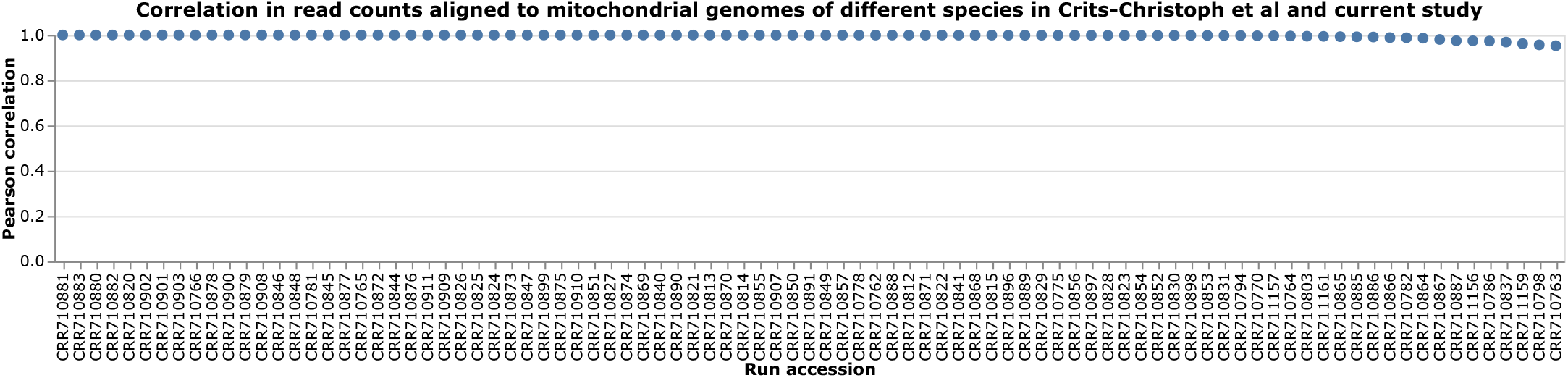
Correlation of the number of read counts mapping to mitochondrial genomes of mammalian species between the current study and CritsChristoph *et al*. (2023) for each sequencing run shared across the studies. The correlation is calculated only across species for which Crits-Christoph *et al*. (2023) provide read counts in the third supplementary figure of their report, and only for runs for which both studies find ≥50 reads mapping to that set of species. See https://jbloom.github.io/Huanan_market_samples/crits_christoph_vs_current_run_corr.html for an interactive version of this plot that enables mouseover of points for details on specific runs, and adjustment of the threshold for how many total aligned mitochondrial reads a run must contain to be shown in this plot.

Despite the strong correlation between my analysis and that by Crits-Christoph *et al*. (2023) on a common set of mammalian mitochondrial genomes, it is important to emphasize that the results are dependent on the genomes to which the reads are aligned. For instance, Figure 2 shows the composition of sample Q61 as determined by aligning reads to different genome sets. My analysis recapitulates the finding of Crits-Christoph *et al*. (2023) that raccoon dogs make the largest contribution to the mammalian mitochondrial genetic material in this sample—but if the analysis is expanded to all chordates, then duck mitochondrial genetic material is more abundant (Figure 2A). But raccoon dog is more abundant than duck if the analysis is instead performed by assembling contigs and then aligning to the full genomes of raccoon dogs, ducks, and several other species, in line with the findings of Crits-Christoph *et al*. (2023). These results are a reminder that the measured metagenomic compo-sition is contingent upon the method and reference set used. Throughout the rest of this paper, I use compositions calculated from reads mapping to mitochondrial genomes rather than contigs mapped to the full genomes for two reasons: (1) this parallels the approach of Crits-Christoph *et al*. (2023) who only reported mitochondrial composition for samples other than Q61, and (2) a number of relevant species only have mitochondrial but not full genomes available in NCBI.

**Figure 2.**
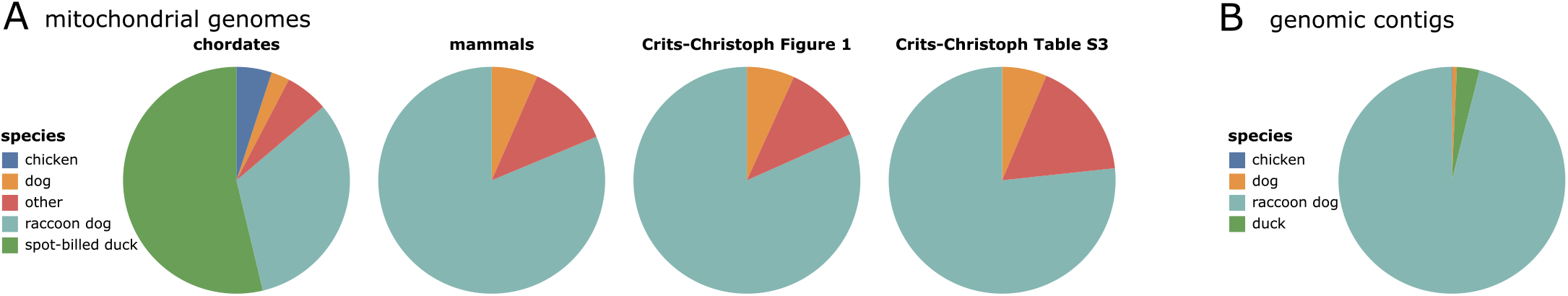
Metagenomic composition of sample Q61. **(A)** Composition as determined by aligning reads to mitochondrial genomes. From left to right: composition determined in current study across all chordates, composition determined in current study across all mammals, composition reported in Crits-Christoph *et al*. (2023) for just mammals shown in the first figure of their report, and composition reported Crits-Christoph *et al*. (2023) across all mammals in the third supplementary table of their report. **(B)** Composition determined in the current study by aligning assembled contigs to the four indicated genomes; this composition is similar to that reported in the third figure of Crits-Christoph *et al*. (2023). See https://jbloom.github.io/Huanan_market_samples/mito_composition.html for an interactive version of (A) that allows similar pie charts to be viewed for any sample. See https://jbloom.github.io/Huanan_market_samples/genomic_contig_composition.html for an interactive version of (B).

The interactive figure at https://jbloom.github.io/Huanan_market_samples/mito_composition.html enables the reader to examine the mitochondrial composition of each sample over both mammals and chordates.

### Most environmental samples contain little or no SARS-CoV-2 reads

To quantify the SARS-CoV-2 content of the samples, I plotted the percentage of all high-quality reads that aligned to SARS-CoV-2 (Figure 3, Table S6, Table S7). Over three-quarters of the samples had no reads aligning to SARS-CoV-2, and most of the rest only had a small number of viral reads.

**Figure 3.**
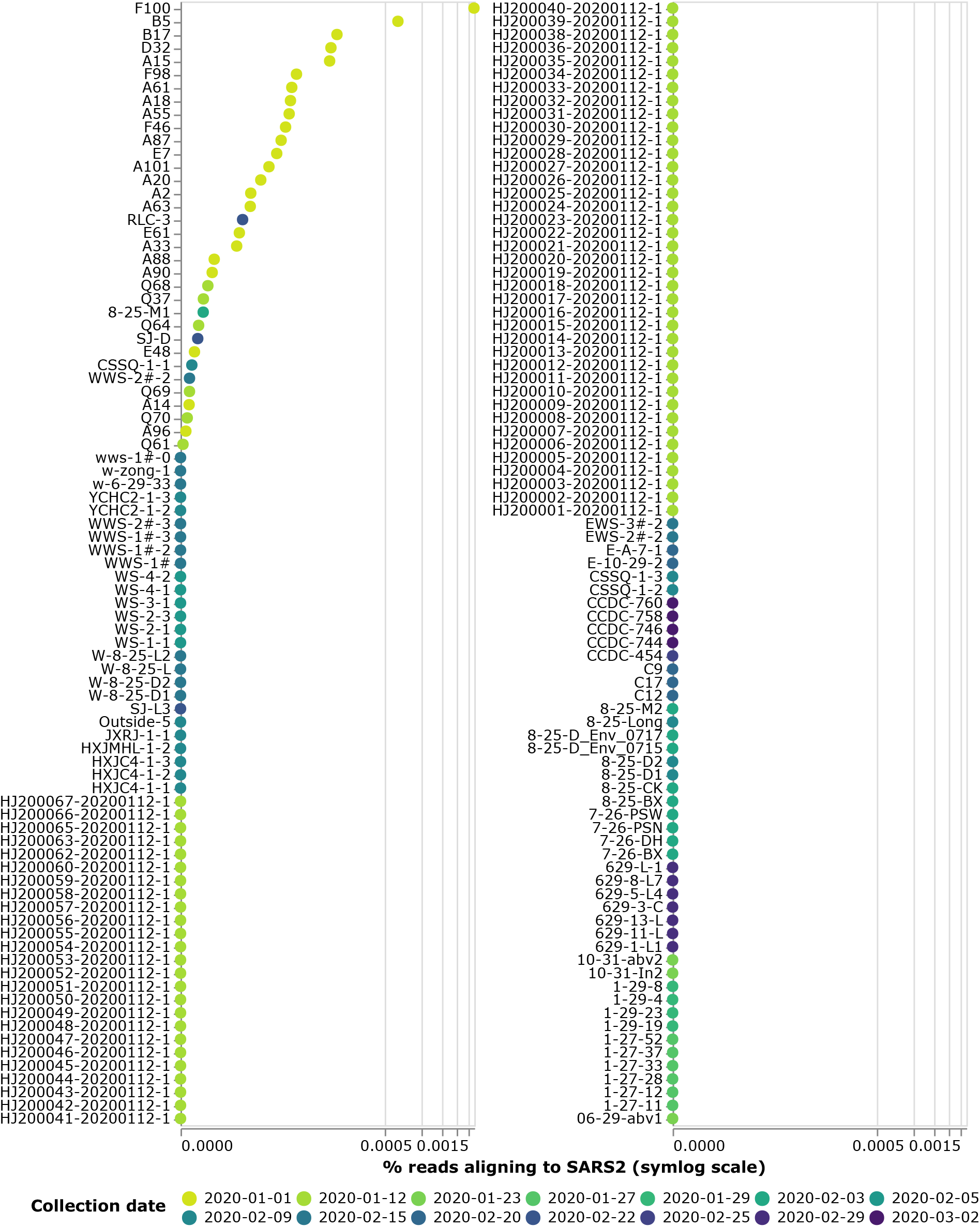
Percentage of high-quality reads that align to SARS-CoV-2 for each sample. Points are colored according to the date that the sample was collected. Note the x-axis uses a symlog scale. See https://jbloom.github.io/Huanan_market_samples/sars2_aligned_vertical.html for an interactive version of this plot where you can mouseover points for details including the mitochondrial composition of each sample, and select only samples from specific dates or from locations.

Despite the fact that most samples were sequenced to depths that exceeded 100,000,000 reads, only two samples had >1,000 reads mapping to SARS-CoV-2 (F100 and B5). These samples were both collected on January-1-2020 and had a chordate mitochondrial composition dominated by catfish and largemouth bass (see interactive version of Figure 3 at https://jbloom.github.io/Huanan_market_samples/sars2_aligned_vertical.html).

Most samples with the highest SARS-CoV-2 content were collected on the Chinese CDC’s first sampling date of January-1-2020, but some samples with SARS-CoV-2 reads were collected on later dates (Figure 3). The non-January-1 sample with the most SARS-CoV-2 was RLC-3, which was collected on February2-2022 and had chordate mitochondrial composition dominated by rat snake, spotted dove, and human (interactive version of Figure 3).

The Q61 sample that was the focus of the raccoon-dog centered media coverage (Wu 2023; Mueller 2023) preceding the report by Crits-Christoph *et al*. (2023) is one of 70 samples collected on January-12-2020 from the west wing of the market. Six of these samples contained SARS-CoV-2 reads, while the other 64 had no SARS-CoV-2 reads (Figure 3). Sample Q61, which has a mitochondrial metagenomic composition dominated by duck and raccoon dog, had 1 of ∼ 2.1 ×10^8^ high-quality reads mapping to SARS-CoV-2 (interactive version of Figure 3). The samples from January-12-2020 with the most SARS-CoV-2 reads were Q68, Q37, and Q64; species that contributed ≥10% of chordate mitochondrial reads in these samples were chicken, dog, duck, Chinese salamander, rabbit, and various snakes.

Overall, there were SARS-CoV-2 reads in just 3 of the 28 samples with at least 20% of their chordate mitochondrial composition from the non-human susceptible species thought to have been sold live in the market (Table 1). There were 14 samples with at least 20% chordate mitochondrial composition from raccoon dogs, but only sample Q61 contained any SARS-CoV-2 reads (1 of ∼2.1 × 10^8^ high-quality reads). None of the six samples with at least 20% chordate mitochondrial composition from bamboo rats contained any SARS-CoV-2 reads. There was one sample each with at least 20% chordate mitochondrial composition from Amur hedgehog and Malayan porcupine that contained SARS-CoV-2 reads (Table 1). If mitochondrial composition is instead analyzed only among mammals rather than chordates, Q61 is still the only sample with at least 20% raccoon dog composition that contains any SARS-CoV-2 reads (Table S8). There is one sample with at least 20% of its mammalian (but not chordate) mitochondrial composition from bamboo rat that contains a small number of SARS-CoV-2 reads (Table S8).

**Table 1.**
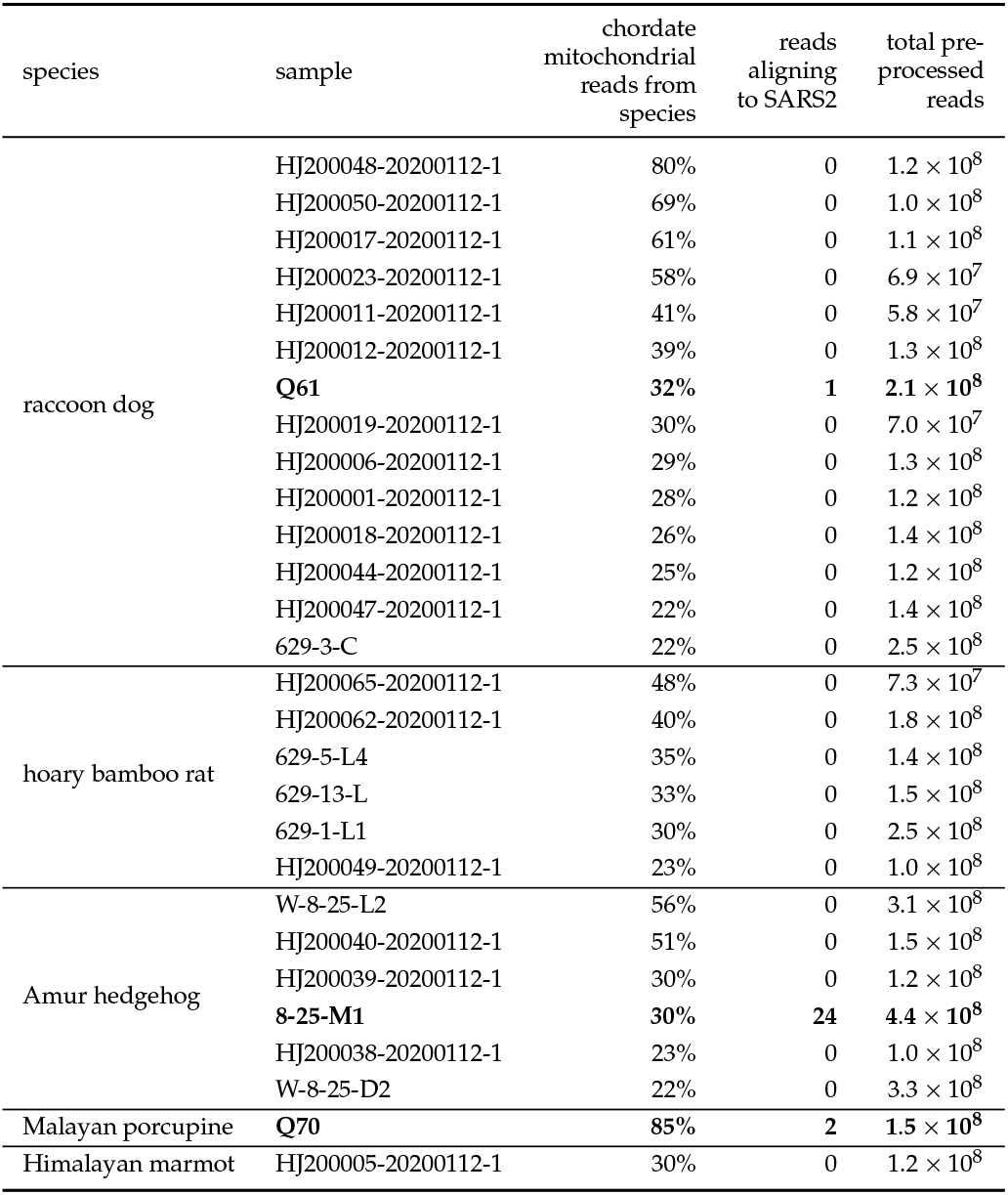
Reads mapping to SARS-CoV-2 out of all high-quality (preprocessed) reads for samples with ≥20% of their chordate mitochondrial composition from a susceptible non-human species as defined in Crits-Christoph *et al*. (2023). Samples with non-zero SARS-CoV-2 reads are in bold. See Table S8 for a similar table that shows mammalian rather than chordate composition. This table uses a 20% cutoff due to space considerations; to see similar data tabulated for all samples with no cutoff, see the much larger Table S9 (for raccoon dog) and Table S5 (for all species).

Note that the 20% cutoff applied in Tables 1 and S8 is not integral to the analysis, but is just a way to subset on samples containing the most material from the species of interest to make the tables small enough to easily visualize. See Tables S9 and S5 for much larger tables that show comparable data for all samples with respect to the raccoon dog or overall mitochondrial composition, respectively. For instance, Table S9 shows that there are a few samples with lower raccoon dog mitochondrial composition that also contain some SARS-CoV-2 reads. However, ultimately these very large tables are difficult to visualize, and so the relationship between SARS-CoV-2 and mitochondrial content for each species for all samples is probably more facilely visualized using the interactive versions of the scatter plots described in the next subsection.

### Correlations of abundance of SARS-CoV-2 to mitochondrial genetic material from various species

To more systematically examine the relationship between SARSCoV-2 and genetic material from different chordates, I calculated the correlation between the number of reads mapping to SARS-CoV-2 versus the mitochondrial genome of each species across all samples. If the correlation is calculated on a log-log scale as was done in the original Chinese CDC preprint (Liu et al. 2022), then the five species whose genetic material is most correlated with the abundance of SARS-CoV-2 are (in order) largemouth bass, catfish, cow, carp, and snakehead fish (Figure 4). None of these species are likely hosts for SARS-CoV-2: non-mammals are not thought to be infectable, and cows were probably sold as animal products rather than live animals. There is a modest correlation between the abundance of SARS-CoV-2 and human mitochondrial material, but this correlation is weaker than for several other species (Figure 4). There is a negative correlation between the abundance of SARS-CoV-2 and mitochondrial material from raccoon dogs and hoary bamboo rats, and at most a weak positive correlation for other susceptible species thought to have been sold live at the market (Figure 4).

**Figure 4.**
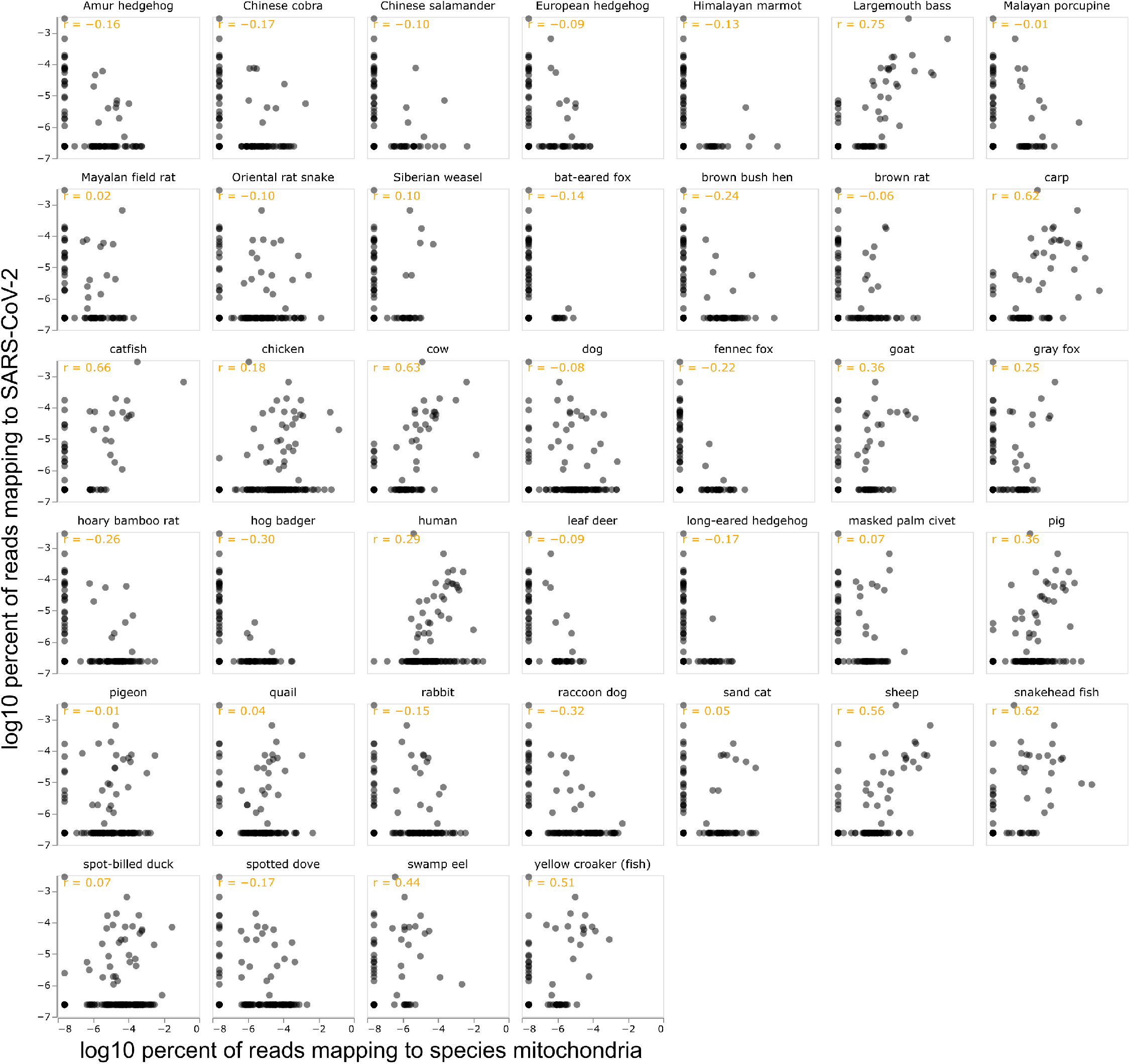
Correlation between percent of all reads mapping to SARS-CoV-2 and the mitochondrial genome of each of the indicated species. Each point represents a different environmental sample, and the orange text shows the Pearson correlation. The scales are log10, and values of zero (which cannot be plotted on a log scale) are shown as half the minimum non-zero value observed across all samples. See https://jbloom.github.io/Huanan_market_samples/per_species_corr_faceted.html for an interactive version of this plot that enables mouseover of points for sample details, selection only of samples collected on specific dates or containing at least one SARS-CoV-2 reads, adjustment of scales from log to linear, and adjustment of mitochondrial percent to be of reads mapping to any mitochondria rather than of all reads. See https://jbloom.github.io/Huanan_market_samples/per_species_corr_single.html for similar plots for individual species. The plots shown here include only samples with at least 200 aligned mitochondrial reads; that option can be adjusted in the interactive plots.

Furthermore, the correlations are highly contingent on the sample set and details of how the statistics are calculated. For instance, if we exclude the first sampling timepoint and only look at samples calculated on January-12-2020 or later, then the most correlated species is the Oriental rat snake, with the correlation for humans becoming even weaker and that for raccoon dogs remaining negative (Figure S1). The correlations also change if only consider samples containing a non-zero number of SARS-CoV-2 reads, or calculate correlations on a linear rather than log scale (Figure S2). The interactive versions of the correlation plots at https://jbloom.github.io/Huanan_market_samples/per_species_corr_faceted.html and https://jbloom.github.io/Huanan_market_samples/per_species_corr_single.html enable the reader to explore the correlations for different sample subsets and methods of calculating the statistics.

### Fully annotated correlation figure analogous to that in the original Chinese CDC preprint

The original Chinese CDC preprint from 2022 had in its fourth figure a plot showing the correlation between the amount of SARS-CoV-2 and genetic material from various animal species in the environmental samples (Liu et al. 2022). The preprint annotated one point in the plot to indicate there was a correlation between the amount of SARS-CoV-2 and human genetic material. However, the preprint failed to annotate the identity of any other species with genetic material that correlated with SARS-CoV-2 abundance. This omission of full species annotations in the correlation plot was widely noted, including in two news articles in *Science* in 2022 (Cohen 2022a,b), and a third article in 2023 that re-printed the incompletely annotated plot (Cohen 2023b). However, neither of the two more recent studies of the metagenomic data have addressed this omission: Crits-Christoph *et al*. (2023) did not report any analysis of the abundance of SARS-CoV-2 reads, and the published 2023 version of the Chinese CDC preprint simply dropped the plot altogether (Liu et al. 2023a).

To remedy this omission, I used the chordate mitochondrial compositions calculated in the current study to generate plots analogous to that in the original Chinese CDC preprint. When the correlations are taken across all sampling dates only for samples containing at least one SARS-CoV-2 read (as done in the original Chinese CDC preprint), then the genetic material of several non-human species (largemouth bass, catfish, cow, sheep, pig) is more correlated with SARS-CoV-2 reads than is human material (Figure 5, upper left). If the plot is expanded to all samples (regardless of whether or not they contain SARS-CoV-2 reads), the trends are broadly similar, with fish remaining the most correlated species (Figure 5, lower left). It is also informative to take the correlations only across samples collected on January-12-2020, which was the date of most intensive sampling of the wildlife stalls (this is when Q61 was collected). For January-12-2020 samples that contain SARS-CoV-2 reads, humans are among the species whose genetic material is most correlated with SARS-CoV-2 reads, but goat and spotted dove have roughly equivalent correlations (Figure 5, upper right). If we include January-12-2020 samples regardless of SARS-CoV-2 content, then snakehead fish and Malayan porcupine are among the most correlated species. Raccoon dog and bamboo rat genetic material are not positively correlated with SARS-CoV-2 reads in any of the sample sets. See the interactive version of Figure 5 at https://jbloom.github.io/Huanan_market_samples/overall_corr.html to explore other sample subsets.

**Figure 5.**
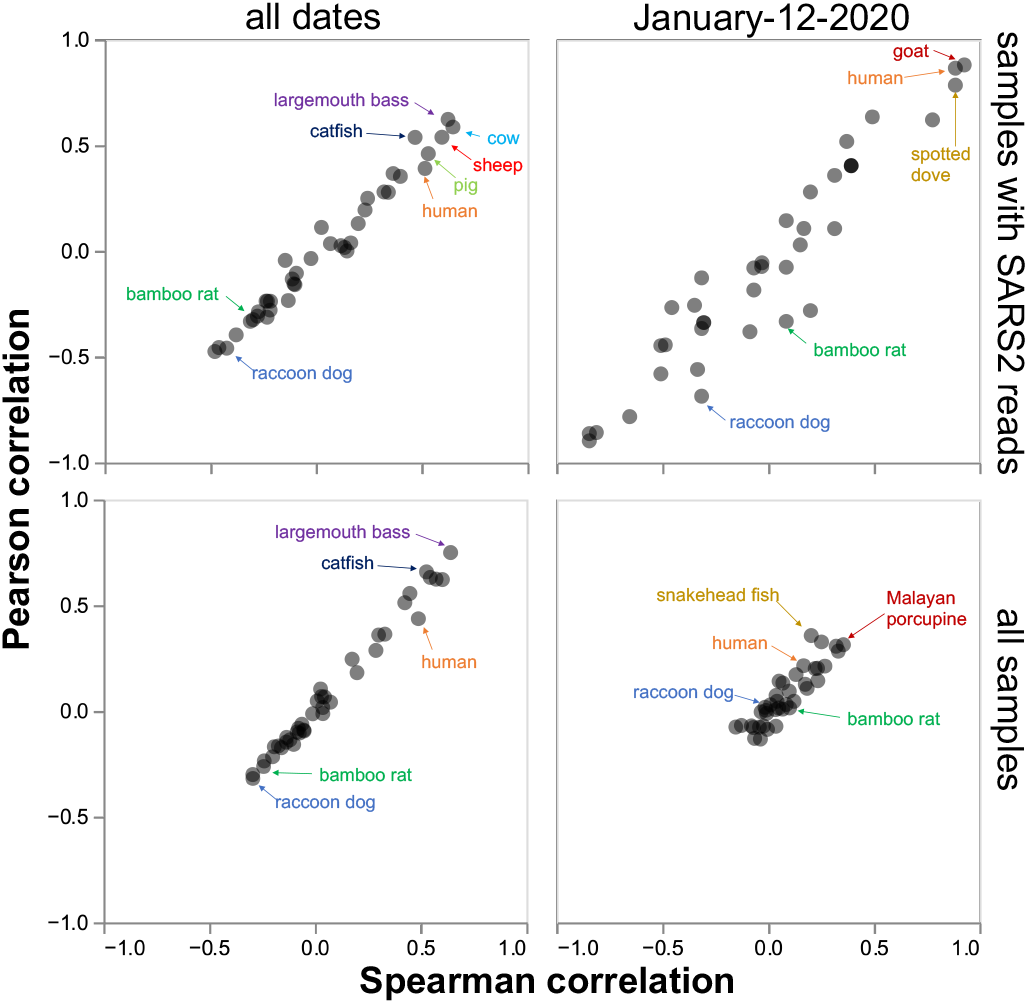
Correlations between SARS-CoV-2 content and mitochondrial content for all species. Top row show just samples containing at least one SARS-CoV-2 read and bottom row shows samples regardless of SARS-CoV-2 content; left shows samples from all sampling dates and right shows just samples from the January-12-2020 date when most of the wildlife sampling occurred. This plot is designed to mimic the fourth figure of Liu *et al*. (2022). Some key species are labeled; see the interactive version at https://jbloom.github.io/Huanan_market_samples/overall_corr.html to mouseover all points for details, select different subsets of samples, and calculate the correlations on a linear or log scale.

Note that the correlation plots in Figure 5 are not exactly identical to those in the original Chinese CDC preprint. Since Liu *et al*. (2022) do not provide sufficiently detailed methods to fully reproduce their analysis, it is impossible to definitively determine the source of the discrepancy—but it is likely due to the inclusion of different genomes in the reference sets used to calculate the metagenomic composition of the samples.

### The low SARS-CoV-2 read counts raise questions about the consistency of the approach use to call sample positivity

The Chinese CDC study includes a table that classifies which environmental samples were positive for SARS-CoV-2 (this is first main table in Liu *et al*. (2022) and the second supplementary table in Liu *et al*. (2023a)). Subsequent studies that have reanalyzed the Chinese CDC data have re-used their classifications of sample positivity (Worobey et al. 2022; Courtier-Orgogozo and de Ribera 2022; Crits-Christoph et al. 2023).

Now that the full data are available, we can examine the criteria used to classify samples as positive or negative for SARS-CoV-2. It appears that the Chinese CDC classified samples as positive if they met *either* of two criteria: they tested positive for SARS-CoV-2 by RT-qPCR, *or* they were metagenomically sequenced and contained at least one read mapping to SARS-CoV-2. However, these criteria are not consistent because not all samples were analyzed by both methods, and the differences in the number of SARS-CoV-2 reads in the sequencing data are often not statistically significant between samples classified as positive versus negative. This inconsistency is illustrated in Table 2 for four example samples: F100 is clearly positive (it both tested positive by RT-qPCR and contained thousands of reads mapping to SARS-CoV-2), but there is no statistical rationale for classifying Q61 as positive but E-10-29-2 and A1 as negative. See Table S10 for merged sequencing and RT-qPCR data for all samples that were either sequenced or called as positive by the Chinese CDC.

**Table 2.** Inconsistency in criteria used to classify SARS-CoV-2 positivity in Chinese CDC study, illustrated with four example samples. There is no consistent rationale for classifying Q61 as positive but E-10-29-2 and A1 as negative: all three were reported negative by RT-qPCR, and A1 was not analyzed by sequencing while the difference in the number of SARS-CoV-2 reads between Q61 and E-10-29-2 is not statistically significant. RT-qPCR results are from the second supplementary table of Liu *et al*. (2023a) (or equivalently the first table of Liu *et al*. (2022)).

| sample | RT-qPCR test result (Ct) | sequencing reads mapping to SARS-CoV-2 out of high-quality reads | classification in Chinese CDC study |
| --- | --- | --- | --- |
| F100 | positive (34.7) | 7,200 out of $2.6 \times 10^8$ | positive |
| Q61 | negative | 1 out of $2.1 \times 10^8$ | positive |
| E-10-29-2 | negative | 0 out of $1.9 \times 10^8$ | negative |
| A1 | negative | not sequenced | negative |

## Discussion

I have described the first full analysis of the association between the abundance of SARS-CoV-2 and genetic material from different animal species in environmental samples collected by the Chinese CDC from the Huanan Seafood Market. This analysis expands upon prior work by Liu *et al*. (2023a) and Crits-Christoph *et al*. (2023) in several ways.

First, I establish that the full set of sequencing data uploaded to public databases by the Chinese CDC at the end of March 2023 contains unmodified versions of all files earlier analyzed by Crits-Christoph *et al*. (2023), as well as a large number of additional files. Therefore, regardless of the disputes about the original access to these files (Cohen 2023a; GISAID 2023; Crits-Christoph et al. 2023), all the files earlier posted on GISAID are now available without alteration in public databases.

Second, I validate the finding of both Crits-Christoph *et al*. (2023) and Liu *et al*. (2023a) that the environmental samples contain genetic material from many species, including humans, various fish, various snakes, cows, goats, pigs, sheep, birds such as ducks and spotted doves, raccoon dogs, bamboo rats, and a long list of other animals. I am only able to quantitatively compare my analyses of metagenomic compositions to the mammalian compositions reported by Crits-Christoph *et al*. (2023), since Liu *et al*. (2023a) provide neither numerical results nor relevant computer code, and Crits-Christoph *et al*. (2023) only provide numerical results for mammalian species. For mammalian species, the compositions determined by my analyses are highly correlated with those reported by Crits-Christoph *et al*. (2023), which provides a reassuring robustness check on both analyses. However, it is important to emphasize that metagenomic compositions depend on the methods and reference genomes used, so future analyses using different approaches (e.g., full versus mitochondrial genomes) would likely yield somewhat different results. I suggest future work should provide full computer code and tabulated results for all species (as done in the current study) to facilitate reproducibility and comparability across studies.

Third, I perform the first comprehensive analysis of the association between the abundance of SARS-CoV-2 and mitochondrial genetic material across all environmental samples. This analysis reveals that the greatest co-mingling of viral and animal material involves species that were almost certainly not infected by SARSCoV-2, such as fish (e.g., largemouth bass, catfish) and livestock (e.g., cows, sheep, goats). Consistent with analyses by the Chinese CDC (Liu et al. 2022), I find some correlation between the abundance of SARS-CoV-2 and human genetic material, but this correlation is weaker than for several non-infectable animals and so on its own is insufficient to identify the source of the viral material. Mitochondrial material from most susceptible nonhuman species sold live at the market is negatively correlated with the presence of SARS-CoV-2: 13 of the 14 samples with at least 20% of their chordate mitochondrial material from raccoon dogs contain no SARS-CoV-2 reads, and the other sample contains just 1 of ∼200,000,000 reads mapping to SARS-CoV-2. Likewise, none of the six samples with at least 20% bamboo rat mitochondrial material contain any SARS-CoV-2 reads. These findings are compatible with the results in Crits-Christoph *et al*. (2023), since that study did not report any analysis of SARSCoV-2 content. However, they are somewhat inconsistent with related media articles that emphasized co-mingling of raccoon dog and viral material (Wu 2023; Mueller 2023)—in fact, raccoon dogs are one of the species with the least co-mingling of their genetic material and SARS-CoV-2. The basic finding that SARSCoV-2 material is not associated with material from susceptible non-human species sold live at the market is largely robust to examining subsets of the samples (such as just those collected on specific dates), and I provide interactive plots to facilitate visualizing the data in different ways.

Fourth, I create a version of the SARS-CoV-2 versus species correlation plot in the fourth figure of the original Chinese CDC preprint (Liu et al. 2022), but with full annotation of all species. The identity of the non-human species correlated with SARS-CoV-2 content had previously been the subject of speculation (Cohen 2022a,b, 2023b). It turns out these species are fish and livestock that are unlikely candidates for having been infected with SARS-CoV-2. Even if the plot is restricted to just samples collected on the date of most intense sampling of wildlife stalls (January-12-2020), no non-human susceptible species is among those most correlated with SARS-CoV-2 content.

Fifth, my analysis calls into question the consistency of the criteria used to classify environmental samples as positive versus negative. For instance, sample Q61 became the subject of widespread media coverage (Wu 2023; Mueller 2023) because it contains raccoon dog genetic material and was classified as SARS-CoV-2 positive by the Chinese CDC. However, this sample tested negative by RT-qPCR and appears to have been called positive on the basis of containing 1 of ∼200,000,000 reads that mapped to SARS-CoV-2. But all environmental samples contain a mix of genetic material from numerous sources, so it is not consistent to classify this particular sample as positive when hundreds of other samples that also tested negative by RT-qPCR are classified as negative because they were never sequenced or had SARS-CoV-2 content statistically indistinguishable from 1 of ∼200,000,000 reads. I suggest that future work analyzing the spatial distribution of SARS-CoV-2 across the Huanan Seafood Market (Worobey et al. 2022; Courtier-Orgogozo and de Ribera 2022; Crits-Christoph et al. 2023) consider the quantitative content of samples (such as determined from deep sequencing in the current study, or the Ct values in Liu *et al*. (2023a)), and only include negative samples subjected to the full set of measurements used to classify other samples as positive.

Overall my study validates the approach of Crits-Christoph *et al*. (2023) of using metagenomic analysis of environmental samples to identify animals and animal products sold at the Huanan Seafood Market. As noted by Crits-Christoph *et al*. (2023), this innovative approach could usefully inform tracing of animals supplied to the market. However, the results described here suggest that the utility of these metagenomic analyses does not extend to indicating whether animals at the market were actually infected by virus. For instance, the presence of 1 in ~200,000,000 sequencing reads mapping to SARS-CoV-2 in a sample containing raccoon dog genetic material does not suggest raccoon dogs were infected, given that material from many other species that certainly were not infected (such as fish) is far more consistently co-mingled with SARS-CoV-2. Of course, the lack of association also does not disprove the possibility of infected animals at the market, particularly at a date substantially preceding the Chinese CDC’s collection of samples—it simply suggests that analysis of the combined viral and animal content of the available environmental samples is not informative for shedding light on this question either way.

When considered in larger context, the inability of the environmental samples to inform on the origins of the virus is unsurprising. These samples were all collected on January-1-2020 or later, which is at least several weeks after the Huanan Seafood Market became a superspreading site for human infections (Li et al. 2020). Therefore, by the time the samples were collected, SARS-CoV-2 had been spread widely across the market by humans regardless of its original source—as evidenced by the results reported here, which show viral genetic material coincident with material from myriad animals ranging from fish to snakes to mammals. The first human infections with SARS-CoV2 in Wuhan occurred no later than November of 2019 (ODNI 2022; Zhang et al. 2020; van Dorp et al. 2020; Pipes et al. 2021), which is over a month before the Chinese CDC reports that it began to collect samples from the market. For this reason, further insight into the origins and early spread of SARS-CoV-2 will likely require learning more about events or cases that occurred no later than November or early December of 2019.

### Limitations of this study

This study has limitations related to the data, methodology, and samples themselves.

For the data, all sequencing files and related annotations derive from information shared by the Chinese CDC. The description of how samples were processed prior to sequencing lacks detail: for instance, Liu *et al*. (2023a) say “human nucleic acid was removed” but not precisely how this was done. Additionally, the data were released after China’s State Council ordered in March of 2020 that all publications and information related to COVID-19 be reviewed by a centralized task force (Kang *et al*. 2020). It is possible that this centralized Chinese government review influenced which data were released (Hvistendahl and Mueller 2023).

For the methodology, the metagenomic compositions were calculated by alignment to a reference set of mitochondrial genomes from a large but still incomplete set of species. It is possible that some species that deposited genetic material do not have exact matches in this reference set, so slightly different results might be obtained if a different reference set was used. Similarly, this study analyzed chordate mitochondrial genomes: different results would be obtained if the reference genome set was expanded (e.g., to all metazoa) or shrunk (e.g., only to mammals), or if the reads were aligned to full rather than mitochondrial genomes. In addition, the results might slightly change with different parameters for read alignment, quality filtering, etc. To ensure transparency of the methodology in this study, I have provided a fully reproducible computational pipeline (https://github.com/jbloom/Huanan_market_samples). In addition, I have provided interactive plots of the results to help the reader explore the effects of different parameter choices on the final results (https://jbloom.github.io/Huanan_market_samples/).

The major limitation of the samples is that they were all collected on January-1-2020 or later, which is well after the first human SARS-CoV-2 infections in Wuhan. The lateness of the sampling relative to the origin of the outbreak limits the conclusions that can be drawn.

## Methods

### Code and data availability

See https://github.com/jbloom/Huanan_market_samples for a GitHub repository containing a fully reproducible Snakemake (Mölder *et al*. 2021) computational pipeline implementing the analysis described in this paper, starting with downloading of the raw data from NGDC and proceeding all the way through to rendering of the plots shown in the figures. That GitHub repository also includes files containing key numerical results. See https://jbloom.github.io/Huanan_market_samples/ for interactive plots of the results rendered using Altair (VanderPlas *et al*. 2018).

### Processing and alignment of deep sequencing data

The FASTQ files with the raw deep sequencing data for all samples were downloaded from the NGDC project CRA010170 (https://ngdc.cncb.ac.cn/gsa/browse/CRA010170). The FASTQ files were pre-processed with `fastp` (Chen et al. 2018) to remove low-quality reads; all tabulations of number of reads in this paper refer to the high-quality reads that passed this pre-processing step.

To create an mitochondrial genome alignment reference set, I followed a procedure partially analogous to that described by Crits-Christoph *et al*. (2023). All mitochondrial genomes in the RefSeq database were downloaded, and then filtered to retain only genomes from the phylum Chordata (in contrast, Crits-Christoph *et al*. (2023) describe retaining all metazoa mitochondrial genomes). The reason I limited to chordate rather than metazoa mitochondrial genomes is that all chordates have very similar length mitochondrial genomes, but at the level of metazoa there is wide variation in mitochondrial genome length. The raccoon dog mitochondrial genome was not present in this set, so it was separately downloaded from Genbank accession KX964606. All the mitochondrial genomes were then filtered to remove highly similar ones. To do this, `Mash` (Ondov et al. 2016) was used to compute the mash distances between all pairs of mitochondrial genomes. To ensure the most relevant mitochondrial genomes were retained, I first specified for manual retention the genomes for all the species listed in the third supplementary file of Crits-Christoph *et al*. (2023), as well as genomes for some additional relevant species (these species are listed under the *mitochondrial_genomes_to_keep* key in https://github.com/jbloom/Huanan_market_samples/blob/main/config.yaml). After retaining all of these manually specified mitochondrial genomes, the pipeline then greedily iterated through all other genomes choosing for retention each genome that was not within a mash distance of 0.07 of another already retained genome. The full set of retained mitochondrial genomes is listed in Table S3.

The retained mitochondrial genomes were then concatenated, and added to the SARS-CoV-2 reference genome (Genbank accession NC_- 045512v2) with its polyA tail trimmed to avoid spurious alignments. This concatenation of the SARS-CoV-2 and mitochondrial genomes served as the reference for all sequencing read alignment.

The sequencing reads were then aligned to this genome using `minimap2` (Li 2018) in the `sr` (short-read) mode. The resulting alignments were filtered to only retain primary alignments with mapping quality of at least 4.

I then used `CoverM` (https://github.com/wwood/CoverM) to compute the number of aligned reads for each run, requiring alignment lengths of at least 40 nucleotides with at least 95% identity, and excluding the 100 nucleotides at the contig ends. The resulting statistics on alignment counts to SARS-CoV-2 and the mitochondrial genomes for each sequencing run are in Tables S4 and S6. To get per-sample alignment counts, I aggregated counts for all runs for each sample to get the statistics in Tables S5 and S7. For the mitochondrial genome compositions and all other analyses in this paper, I only included only the samples from metagenomic sequencing, which are annotated with the description “RNA sequencing of total nucleic acids from environmental swabs for metagenomics” in the metadata provided by Liu *et al*. (2023a). Species were only retained in the annotated reference set if they had at least 20% of all aligned mitochondrial reads and at least 4000 covered based for at least one run, or if they are one of the species specified under the *mitochondrial_genomes_to_keep* key in https://github.com/jbloom/Huanan_market_samples/blob/main/config.yaml. The species specified under that key include all susceptible species studied in Crits-Christoph *et al*. (2023), so this criteria does not lead too dropping of any of the species in their study.

### Assembling and aligning contigs for sample Q61

For sample Q61, contigs were also assembled and aligned to full genomes to facilitate comparison to Crits-Christoph *et al*. (2023), who also performed a similar analysis only for this sample. The contigs were assembled using `Trinity` (Grabherr et al. 2011), and then aligned to using `minimap2` (Li 2018) to concatenated full genomes for chicken, dog, raccoon dog, and duck. Contig alignments were only reported if they had a mapping quality of at least 10, an alignment length of at least 300 nucleotides, and an identity of at least 98%.

## Acknowledgments

I thank Liu *et al*. (2023a) for publicly sharing via the NGDC database the deep sequencing data analyzed in this study. An initial version of this preprint was revised in part based on helpful comments posted on *bioRxiv* by Florence Débarre and Alex Crits-Christoph (see appendix posted as supplementary material). This study utilized the Fred Hutch Scientific Computing infrastructure, which is supported in part by NIH grants S10-OD-020069 and S10-OD-028685. The author is an Investigator of the Howard Hughes Medical Institute.

---

**Figure S1.**
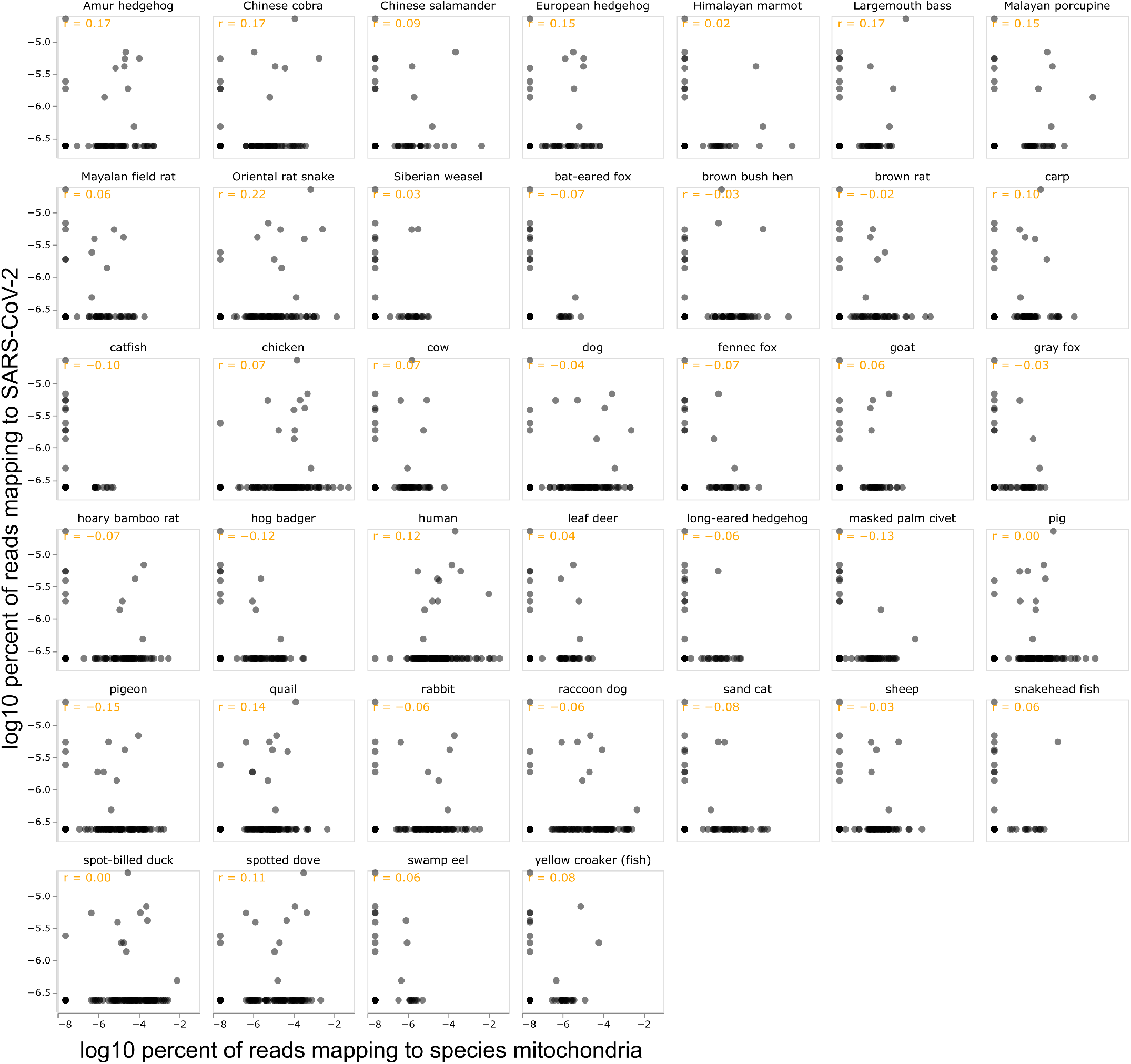
Correlation between percent of all reads mapping to SARS-CoV-2 and the mitochondrial genome of each of the indicated species only among samples collected after the first (Jan-1-2020) sampling date. Except for the fact that this figure excludes samples from the first date, this figure is otherwise the same as Figure 4. See https://jbloom.github.io/Huanan_market_samples/per_species_corr_faceted.html for an interactive version of this plot with numerous options, including allowing further refinement of the sample date range. The plots shown here include only samples with at least 200 aligned mitochondrial reads; that option can be adjusted in the interactive plots.

**Figure S2.**
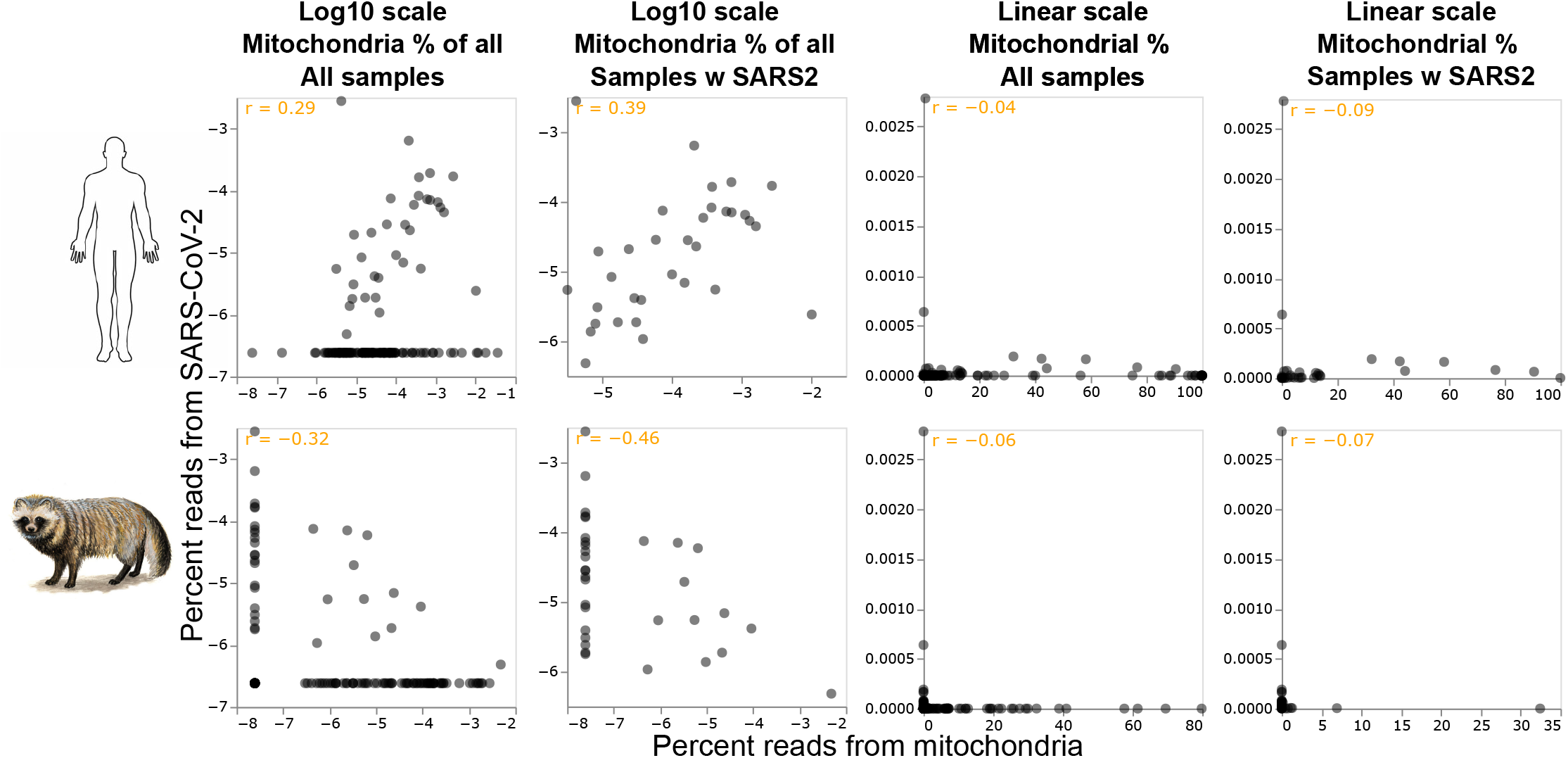
The correlations between the amount of SARS-CoV-2 and mitochondrial genetic material in the samples depends on how the correlation is calculated. The top row shows correlations for human, and the bottom row shows correlations for raccoon dog. Each column shows the correlations calculated different ways: with a log or linear scale, calculating the mitochondrial percent to be of all reads or just reads mapping to the mitochondria, and showing all samples or only those with at least one SARS-CoV-2 read. See https://jbloom.github.io/Huanan_market_samples/per_species_corr_single.html for interactive versions of the plot panels. The plots shown here include only samples with at least 200 aligned mitochondrial reads; that option can be adjusted in the interactive plots.

**Table S1** See https://github.com/jbloom/Huanan_market_samples/blob/main/results/crits_christoph_data/check_sha512_vs_crits_christoph.csv for the correspondence between the FASTQ files analyzed by Crits-Christoph *et al*. (2023) and the files deposited in the NGDC by Liu *et al*. (2023a). The files are matched based on their SHA-526 hashes, which were computed directly for the NGDC files as part of the current study, and were taken from Table S1 of Crits-Christoph *et al*. (2023) for that study.

**Table S2** See https://github.com/jbloom/Huanan_market_samples/blob/main/results/metadata/merged_metadata.csv for the metadata for all of the samples and files uploaded to the NGDC by Liu *et al*. (2023a) under accession CRA010170.

**Table S3** See https://github.com/jbloom/Huanan_market_samples/blob/main/results/mitochondrial_genomes/retained.csv for the NCBI accessions of the set of chordate mitochondrial genomes that was used as the alignment reference for the metagenomic analysis.

**Table S4** See https://github.com/jbloom/Huanan_market_samples/blob/main/results/aggregated_counts/mito_composition_by_run.csv for the counts of reads aligning to each mitochondrial genome for each sequencing run.

**Table S5** See https://github.com/jbloom/Huanan_market_samples/blob/main/results/aggregated_counts/mito_composition_by_sample.csv for the counts of reads aligning to each mitochondrial genome for each sample.

**Table S6** See https://github.com/jbloom/Huanan_market_samples/blob/main/results/aggregated_counts/sars2_aligned_by_run.csv for the counts of reads aligning to SARS-CoV-2 for each sequencing run.

**Table S7** See https://github.com/jbloom/Huanan_market_samples/blob/main/results/aggregated_counts/sars2_aligned_by_sample. csv for the counts of reads aligning to SARS-CoV-2 for each sequencing sample.

**Table S8.** Reads mapping to SARS-CoV-2 out of all high-quality (pre-processed) reads for samples with ≥20% of their mammalian mitochondrial composition from a susceptible non-human species as defined in Crits-Christoph *et al*. (2023). Samples with non-zero SARS-CoV-2 reads are in bold. See Table 1 for a similar table that shows chordate rather than mammalian composition. This table uses a 20% cutoff due to space considerations; to see similar data tabulated for all samples with no cutoff, see the much larger Table S9 (for raccoon dog) and Table S5 (for all species).

| species | sample | mammalian<br>mitochondrial<br>reads from<br>species | reads<br>aligning<br>to SARS2 | total pre-processed<br>reads |
| --- | --- | --- | --- | --- |
| raccoon dog | HJ200050-20200112-1 | 94% | 0 | 1e+08 |
|  | HJ200023-20200112-1 | 93% | 0 | 6.9e+07 |
|  | HJ200048-20200112-1 | 92% | 0 | 1.2e+08 |
|  | HJ200019-20200112-1 | 90% | 0 | 7e+07 |
|  | **Q61** | **81%** | **1** | **2.1e+08** |
|  | HJ200017-20200112-1 | 81% | 0 | 1.1e+08 |
|  | 629-3-C | 78% | 0 | 2.5e+08 |
|  | HJ200044-20200112-1 | 76% | 0 | 1.2e+08 |
|  | HJ200014-20200112-1 | 70% | 0 | 1.1e+08 |
|  | HJ200047-20200112-1 | 66% | 0 | 1.4e+08 |
|  | HJ200011-20200112-1 | 61% | 0 | 5.8e+07 |
|  | HJ200018-20200112-1 | 53% | 0 | 1.4e+08 |
|  | HJ200013-20200112-1 | 50% | 0 | 1.1e+08 |
|  | HJ200012-20200112-1 | 50% | 0 | 1.3e+08 |
|  | HJ200043-20200112-1 | 48% | 0 | 8.3e+07 |
|  | HJ200027-20200112-1 | 43% | 0 | 9e+07 |
|  | HJ200024-20200112-1 | 42% | 0 | 9.8e+07 |
|  | HJ200001-20200112-1 | 38% | 0 | 1.2e+08 |
|  | HJ200016-20200112-1 | 33% | 0 | 1.2e+08 |
|  | HJ200007-20200112-1 | 31% | 0 | 1.4e+08 |
|  | HJ200020-20200112-1 | 31% | 0 | 7.9e+07 |
|  | HJ200006-20200112-1 | 30% | 0 | 1.3e+08 |
|  | HJ200009-20200112-1 | 27% | 0 | 1.5e+08 |
|  | HJ200045-20200112-1 | 27% | 0 | 1.1e+08 |
|  | HJ200026-20200112-1 | 25% | 0 | 1e+08 |
|  | HJ200022-20200112-1 | 23% | 0 | 1.2e+08 |
|  | HJ200004-20200112-1 | 23% | 0 | 2.3e+08 |
|  | HJ200065-20200112-1 | 22% | 0 | 7.3e+07 |
| hoary bamboo rat | 629-13-L | 85% | 0 | 1.5e+08 |
|  | HJ200041-20200112-1 | 56% | 0 | 1.8e+08 |
|  | HJ200015-20200112-1 | 56% | 0 | 1.4e+08 |
|  | HJ200065-20200112-1 | 56% | 0 | 7.3e+07 |
|  | 629-1-L1 | 55% | 0 | 2.5e+08 |
|  | 629-5-L4 | 54% | 0 | 1.4e+08 |
|  | HJ200062-20200112-1 | 42% | 0 | 1.8e+08 |
|  | HJ200013-20200112-1 | 37% | 0 | 1.1e+08 |
|  | HJ200049-20200112-1 | 31% | 0 | 1e+08 |
|  | HJ200046-20200112-1 | 28% | 0 | 1.1e+08 |
|  | HJ200030-20200112-1 | 28% | 0 | 1.1e+08 |
|  | HJ200067-20200112-1 | 27% | 0 | 7.7e+07 |
|  | HJ200025-20200112-1 | 21% | 0 | 1.2e+08 |
|  | **Q68** | **20%** | **6** | **8.7e+07** |
| Amur hedgehog | W-8-25-L2 | 76% | 0 | 3.1e+08 |
|  | HJ200040-20200112-1 | 75% | 0 | 1.5e+08 |
|  | HJ200038-20200112-1 | 63% | 0 | 1e+08 |
|  | HJ200039-20200112-1 | 62% | 0 | 1.2e+08 |
|  | 8-25-CK | 62% | 0 | 1.5e+09 |
|  | W-8-25-L | 59% | 0 | 2.9e+08 |
|  | HJ200035-20200112-1 | 59% | 0 | 7.2e+07 |
|  | **8-25-M1** | **53%** | **24** | **4.4e+08** |
|  | HJ200033-20200112-1 | 51% | 0 | 1.2e+08 |
|  | W-8-25-D2 | 50% | 0 | 3.3e+08 |
|  | W-8-25-D1 | 50% | 0 | 2.9e+08 |
|  | HJ200045-20200112-1 | 41% | 0 | 1.1e+08 |
|  | HJ200032-20200112-1 | 35% | 0 | 1.4e+08 |
|  | 8-25-M2 | 24% | 0 | 1.6e+09 |
|  | HJ200036-20200112-1 | 21% | 0 | 2.2e+08 |
| Malayan porcupine | **Q70** | **93%** | **2** | **1.5e+08** |
|  | HJ200001-20200112-1 | 23% | 0 | 1.2e+08 |
| Himalayan marmot | HJ200005-20200112-1 | 36% | 0 | 1.2e+08 |
| masked palm civet | HJ200042-20200112-1 | 22% | 0 | 1.9e+08 |

**Table S9** See https://github.com/jbloom/Huanan_market_samples/blob/main/results/plots/raccoon_dog_long.csv for a table giving the raccoon dog mitochondrial composition (both among chordates and mammals) along with the SARS-CoV-2 content for all samples.

**Table S10** See https://github.com/jbloom/Huanan_market_samples/blob/main/results/rt_qpcr/rt_qpcr.csv for combined RT-qPCR data and number of SARS-CoV-2 reads for all environmental samples that were either RT-qPCR positive or metagenomically sequenced. This table was constructed by merging the results for all sequenced samples with all samples listed in the second supplementary table of Liu *et al*. (2023a) (or equivalently the first main table of Liu *et al*. (2022)), which lists all “positive samples.” Since the Chinese CDC says all environmental samples were tested by RT-qPCR, samples not listed as RT-qPCR positive are presumed to have tested negative by that assay. Note that this table does *not* list samples that tested negative by RT-qPCR and were not sequenced.

