## Supplementary Appendix for "Association between SARS-CoV-2 and metagenomic content of samples from the Huanan Seafood Market"

### Appendix: response to comments and revision of first version of preprint

After the original version of this preprint was posted on *bioRxiv* at https://www.biorxiv.org/content/10.1101/2023.04.25.538336v1, Florence Débarre and Alex Crits-Christoph posted a set of helpful comments at https://www.biorxiv.org/content/10.1101/2023.04.25.538336v1#comments using the *bioRxiv* comment feature.

This appendix provides a response to those comments, some of which were used to inform revisions incorporated into this new version of the preprint. Note that the computer code / data / paper from the initial version are all also available on GitHub under tag `bioRxiv_v1` (https://github.com/jbloom/Huanan_market_samples/tree/bioRxiv_v1) and the code / data / paper for this revised version are under the tag `bioRxiv_v2` (https://github.com/jbloom/Huanan_market_samples/tree/bioRxiv_v2). To see all differences between the two versions, go to https://github.com/jbloom/Huanan_market_samples/compare/bioRxiv_v1...bioRxiv_v2 which shows the differences between the GitHub tags.

In addition to the revisions made in response to the comments below, I also made a minor tweak to ensure that the same `fastp` pre-processing filters were being applied to both the single- and paired-end sequencing data. Overall, none of the modifications to the code led to anything more than minor changes in the overall results.

Below the comments posted on *bioRxiv* by Florence Débarre and Alex Crits-Christoph are in **purple text**, and my responses are in **black text**.

#### *Comments and responses*

In this preprint, Bloom re-analyses a dataset of metagenomic data recently shared on open repositories by Liu *et al.* (2023a). A subset of these data, previously made available on GISAID, was also previously analyzed by Crits-Christoph *et al.* (2023)—of which we are co-authors.

1. Bloom's analysis largely confirms the genetic identification of wildlife species by Crits-Christoph *et al.* (2023).

   The identification of animal species differed between Liu *et al.* (2023a) and Crits-Christoph *et al.* (2023), in particular regarding the abundance of raccoon dog genetic material in sample Q61. Bloom's analysis, done with similar but independent methods from Crits-Christoph *et al.* (2023), largely replicates their findings.

   A comprehensive Github repository allows any interested reader to conveniently check Bloom's results by themselves.

   I agree that my analysis largely confirm the analysis of the mammalian mitochondrial genetic material in your study. As I note in my preprint, it is reassuring to have two independent studies reach largely consistent results on this point. As I also note, I think the main value of your earlier work was to provide genetic confirmation of which animals were at the market, and my study basically independently confirms your findings on that point.

   I am glad you find the GitHub repository useful!

2. The correlation analysis presented by Bloom contains numerous flaws.

   Bloom reproduced the correlation analysis initially presented in Liu *et al.* (2022), but Bloom's analysis, as conducted, is inappropriate to answer the question of which animal hosts shed the viral material detected. Crits-Christoph *et al.* (2023) were missing data to carry out such an analysis, but warned against its use. Here we repeat and expand the arguments presented in Crits-Christoph *et al.* (2023). These points would need to be taken into account for a correlation analysis to be valid, but were not by Bloom (and often just cannot be):

   At a high level, I agree that the lack of any correlation between SARS-CoV-2 and the genetic material of susceptible animals does not *disprove* the possible existence of an infected animal (for reasons you list below). However, it does indicate that the genetic content of these samples is insufficient to reliably indicate whether any animal was infected. That is the claim my preprint makes. In the end, if the SARS-CoV-2 content is not associated with the genetic content from susceptible animals, then certainly the genetic material cannot reliably indicate that the animals were infected and shedding the virus.

   I think we are in agreement that co-mingling of viral and animal genetic material in these samples is insufficient to indicate if any animals were infected. Regardless of perceptions/discussion on social or popular media, your report never claimed proof of infected animals, and my preprint never claimed disproof of infected animals. Rather, my understanding is that by fully analyzing the data we are converging on the view that the genetic content of these samples is simply insufficient to reliably determine if any animals were infected.

   a) The outbreak had been ongoing for weeks in the market when the samples were collected, and multiple humans had been infected. By then, most virus in the market will come from human cases, especially as stalls containing human cases were targeted first, when sampling was performed on January 1. The large number of samples from these stalls with sick humans will highly influence correlational analyses. As shown in Worobey et al. (2022) on similar data obtained via qPCR, both the distance from known human cases and the distance from wildlife stalls independently contributed to the spatial variation in viral positive samples. The number of human cases at the time of sampling means that even if infected animals were present at the market, an underpowered correlation analysis of the sequencing data would be unlikely to have revealed it.

I agree that the late date of sample collection is a major limitation, and mention that several times in my preprint. As a say in the preprint, by the time the samples were collected humans had spread the virus widely across the market. That means the environmental samples are not particularly informative on the original source of the virus in the market in either direction, which is the central conclusion I draw.

b) The correlation analysis treats all animals of a given species as an homogeneous group. There were however multiple stalls selling the same animal species, but likely not from the same supply sources. The presence of uninfected animals in one stall would affect the result of a correlation analysis, but would not invalidate the potential presence of infected animals in another stall.

   Certainly I agree that this is a limitation. As I say in the preprint, these environmental samples are not sufficient for reliably indicating if any animals were infected.

c) The samples themselves are not homogeneous: for instance, within-sample animal diversity is different when the swab was taken directly from fish packaging than from the ground.

   In addition, wildlife stall samples tend to be composed of a diverse menagerie of animal species RNA/DNA (a consequence of animals housed directly on top of each other and equipment shared between them), while samples related to human cases elsewhere in the market tend to be fairly simple. Therefore, by correlating abundances within just the positive samples, human-shed samples tend to have very high relative abundances for human DNA (or, e.g., human and fish), while animal shed samples tend to have lower abundances for any given species. This is a natural consequence of sequencing abundances being relative, not absolute.

   There is no correlation between the amount of SARS-CoV-2 genetic material and that from potentially susceptible animals like raccoon dogs or bamboo rats regardless of whether the correlation is taken over just SARS-CoV-2 positive samples or all samples. This is shown in the new Figure 5 and can also be seen in Figure 4 and perhaps most easily in the interactive version of that figure linked in its legend.

d) SARS-CoV-2 read counts can differ for multiple reasons: there can be different quantities of virus shed in the first place, and different times since virus deposition will influence detection, as RNA degrades over time. While it is not possible to precisely date the time of viral deposition, there are two important considerations. First, if wildlife introduced the virus first to the market, animal-shed virus would (on average) occur before human sheddings. Second, wildlife stalls were sampled multiple days after stalls with reported cases (January 12 vs January 1). Put together, this means there would have been substantially more time for virus shed by animals to decay than virus shed by humans.

   There is no correlation between the amount of SARS-CoV-2 and genetic material from potentially susceptible animals like raccoon dogs or bamboo rats regardless of whether the analysis is performed on all samples or restricted to just samples collected on later dates (such as January 12). This can be seen in Figure 5, Figure S1, and perhaps most easily in the interactive plots at https://jbloom.github.io/Huanan_market_samples/per_species_corr_faceted.html, which enable subsetting to samples collected on arbitrary dates. Of course this fact does not rule out the possibility that animals shed virus earlier and that viral RNA decayed. It is also possible animals never shed virus. Both possibilities are consistent with my conclusion that these samples are not sufficient to reliably indicate whether any animals were infected.

e) Virus presence is detected through RNA, host presence through mtDNA, which degrade differently over time. Relative abundances therefore depend on the time difference between when the different genetic materials were deposited and when they were collected.

   See response to prior point.

f) Stalls were sampled multiple times. A correlation analysis would need to take into account spatial proximity and in particular model the per-stall effect.

   At a qualitative level (just for instance looking at the scatter plots), it is obvious that the samples that contain substantial amounts of material from potentially susceptible animals like raccoon dogs or bamboo rats contain no or very little SARS-CoV-2 (see interactive plots at https://jbloom.github.io/Huanan_market_samples/per_species_corr_single.html). This is going to remain true no matter how the samples are partitioned or modeled. Given this fact, it seems clear that the samples do not suggest shedding by an infected animal. Of course there could have been infected animals earlier or that shed virus that was not collected in these samples, but in either case that makes these samples uninformative for determining if any animal was infected.

g) The correlations reported in Bloom's figure 5 are calculated solely on samples that contained any positive reads for SARS-CoV-2, and were therefore sequencing-positive for the virus. Therefore, this is not a test of which species were more likely to be in samples that tested positive for the virus, but how abundant each species was in the samples that did test positive. This is an important distinction.

The correlations in Figure 4 and its interactive version (https://jbloom.github.io/Huanan_market_samples/per_species_corr_faceted.html) show all samples, and I have also added panels with all samples to Figure 5. It remains the case even when looking at all samples that SARS-CoV-2 content is mostly associated with material from animals that definitely were not infected.

For these reasons, both positive and negative correlations are largely uninterpretable:

- A positive correlation is not indicative of infection. As an enlightening example (also highlighted in Crits-Christoph *et al.* (2023)), the most correlated species in Bloom's analysis is a fish species that is not susceptible to SARS-CoV-2.
- A negative correlation is not indicative of lack of infection (see above).

I agree with this essential point: my preprint simply concludes that the genetic content of these samples is insufficient to reliably determine if there were any infected animals.

3. Some samples appear to be missing in the analysis, which contained high viral read numbers: A20, F13, F54.

   In my analysis, I filter only for samples sequenced metagenomically, and exclude samples that were enriched for virus for viral sequencing. Originally, I was filtering all samples that had any virus-enrichment based sequencing. In the updated pipeline, for samples like A20 that have both metagenomic and viral sequencing, I now retain the metagenomic sequencing while ignoring the viral sequencing—thanks for pointing out this oversight. This adds a few samples including A20, but does not change any of the overall results. However, F13 and F54 are still excluded because according to the metadata in the *Experiments* sheet in the Chinese CDC metadata Excel file at https://ngdc.cncb.ac.cn/gsa/browse/CRA010170, these samples were not sequenced metagenomically (there are no entries with the "RNA sequencing of total nucleic acids from environmental swabs for metagenomics" annotation for them in the *Experiments* sheet).

4. Bloom chose to align reads to chordates' mitochondrial genomes, but did not justify this choice. Only mammals are known to be infected by SARS-CoV-2, so an analysis of just mammalian RNA/DNA, as presented in Crits-Christoph *et al.* (2023), is most relevant to the question of which species may have shed the virus in a particular sample. If the aim was to provide a better picture of the diversity of animals sold at the Huanan market, all metazoa should have been included, to be able to identify seafood for instance (a key early case was a shrimp seller).

   However, the end result is that Bloom's 20% chordate inclusion threshold for this table results in most of the samples with both wildlife RNA/DNA (e.g. raccoon dogs) and viral RNA being excluded. The claim that only one sample of all samples with raccoon dog DNA that contains SARS-CoV-2 is therefore inaccurate: please see the supplementary data of either Crits-Christoph *et al.* (2023) or Bloom et al. for comparative results on the true number of such samples (which is at least 5, depending on the given analysis).

   The reason I limited to chordate rather than metazoa mitochondrial genomes is that all chordates have very similar length mitochondrial genomes, but at the level of metazoa there is wide variation in mitochondrial genome length—this is now explained in the revised Methods.

   In any case, the choice of using chordate versus mammalian has no real effect. The 20% cutoff in Table 1 was just to make the table small enough to easily fit in the main text. I have now added a comparable table (Table S8) that lists mammalian rather than chordate composition, and another table that lists SARS-CoV-2 and raccoon dog content for all samples without any cutoff (Table S9). The essential result is unchanged: the samples with abundant raccoon dog mitochondrial material have little or no SARS-CoV-2 reads. This is also easily seen in the interactive scatter plots provided in the paper.

   There are a few other samples with low raccoon dog content (less than 20% of either their chordate or mammalian mitochondrial content, see Table S9) that have some SARS-CoV-2 reads, but these samples also have more abundant genetic material from many other species. So certainly they are not sufficient to support an association between viral and raccoon dog material that exceeds that of any of a large number of other species that clearly were not infected (because they are not susceptible). This all supports the conclusion that the genetic contents of these samples are insufficient to conclude if any animal was infected.

5. Bloom's comments on the definition of positive samples overlook the difference in sensitivity between qPCR and sequencing in environmental sampling—qPCR is highly sensitive, with near single molecule detection resolution, and correlates with the absolute load of the virus in the sample. Sequencing, however, is both less sensitive (even when deeply sequencing), and highly dependent on the quantity of what else is in the sample. Thus, it is very challenging to identify SARS-CoV-2 in samples sequenced from the environment without any kind of enrichment performed: this is because the virus is comparatively rare compared to abundant environmental microbes (bacteria, fungi) as well as DNA/RNA shed by animals and humans. This is why most environmental viral sequencing is done after performing a viral enrichment (either physical or amplicon). For a similar comparison of unenriched SARS-CoV-2 sequencing, see the supplementary table

S1 of Crits-Christoph et al. 2021 ("Genome Sequencing of Sewage Detects Regionally Prevalent SARS-CoV-2 Variants"), where 5 unenriched samples with viral Ct values in the range of 30-32 were deeply sequenced, and only 1-17 SARS-CoV-2 reads were obtained. For this reason, we do not encourage using only sequencing data quantitative values and ignoring qPCR positive samples. There is no indication in the Liu *et al.* (2023a) dataset that we have seen to indicate the qPCR results were impacted by contamination. Indeed, 112/116 samples that were reported negative for qPCR were also negative by NGS, indicating a high degree of agreement between the two approaches, and that qPCR is merely more sensitive.

In addition, the definition of a positive sample cannot be done independently of the provenance of the sample: even at very low read count, the presence of other positive samples in the same stall at the same date is indicative of sample positivity.

I agree that RT-qPCR could be more sensitive than sequencing. But the key point is that the "positive" sample with the most raccoon dog genetic material (Q61) tested *negative* by RT-qPCR, and was called positive by the Chinese CDC on the basis of having 1 in 200,000,000 reads align to raccoon dog. It is not consistent to call that sample "positive" if other samples that also tested negative by RT-qPCR but were never sequenced are called "negative." Samples should only be analyzed together if they were subjected to the same consistent set of assays: so for instance, all samples could be classified by RT-qPCR, or all samples subjected to both assays could be classified based on both. This observation is not a criticism of your work: the basis for these classifications only recently became clear with the new Liu *et al.* (2023a) paper. I just think we should now consider these points now that they are apparent.

As far as other samples in the same stall, it is clear that SARS-CoV-2 was spread widely across the market: that is why we see it on samples full of material from fish, snakes, etc. Therefore, it is my contention that we simply can't conclude much about whether any animal was infected if the virus is spread widely in the environment uncorrelated with any specific animal genetic material. This is true regardless of whether we do the analysis at the level of samples or stalls.

6. Citations to news articles are vague or misleading. It would be preferable to stick to arguments presented in scientific documents, rather than rely on newspaper articles.

   a) As rationale for his analysis, in the Introduction of his preprint, Bloom claims that "Other scientists pointed out that if the raw data were shared, it would be possible to expand upon the analysis in the 2022 Chinese CDC preprint to determine if the abundance of SARS-CoV-2 genetic material correlated with material from other species (Cohen 2022a,b)", where the two Cohen citations are news articles. The authors quoted in these articles (Robertson, Rambaut, Holmes) did not mention Figure 4a because they thought the analysis was useful enough to be reproduced, but because it was possible that other animals' genetic material was present in the environmental samples, and these data were not shown.

      The overall rationale of my study was to examine the quantitative association between SARS-CoV-2 content and animal genetic material. Figure 4a of the original Chinese CDC study was the only prior work on this topic, and it was widely discussed including in the three Cohen news articles I mentioned, one of which even re-printed the figure. I think it is important to provide this as background, and I am careful to distinguish what was discussed in Crits-Christoph *et al.* (2023) (which did not discuss this figure) versus news articles.

   b) By conflating their views with misinterpretations of news reports written about their work, Bloom misrepresents the conclusions of Crits-Christoph *et al.* (2023): Crits-Christoph *et al.* (2023) did not conclude they had found the intermediate host and it was raccoon dogs: their report clearly stated that they could not "identify the intermediate animal host species from these data", and that the presence of infected animals was a "plausible explanation" for their results. And they explicitly stated that "the most abundant animal in the sequencing data of a particular sample is not necessarily the source of the virus in that sample".

      I am careful to delineate that Crits-Christoph *et al.* (2023) did *not* analyze the SARS-CoV-2 content in their report, and so never claimed that animal and viral co-mingling in the environmental samples indicated an infected animal. My preprint mentions several times how it is consistent with the results reported in Crits-Christoph *et al.* (2023), but adds more analyses that look at association between animal and viral material.

      However, it is true that the news articles I cite *did* talk about co-mingling of material suggesting infected animals, and those were widely read. I therefore provide the full context to help disentangle the confusion between what was in the scientific report versus the related news articles, which the scientific authors obviously did not write.
